# Molecular Basis of Angicin Activity

**DOI:** 10.64898/2026.06.03.729741

**Authors:** Verena Vogel, Armando A. Rodríguez, Beata Kruszewska-Naczk, Stefanie Mauerer, Lia-Raluca Olari, Janet Köhler, Puja Yadav, Jana Apolloni, Clarissa Read, Paul Walther, Gilbert Weidinger, Jan Münch, Ludger Ständker, Barbara Spellerberg

**Affiliations:** Institute for Medical Microbiology and Hygiene, Ulm University Hospital, Ulm, Germany; ULMTec Core Facility Functional Peptidomics, Ulm University Hospital, Ulm, Germany; ULMTec Core Facility of Mass Spectrometry and Proteomics, Ulm University Hospital, Ulm, Germany; Laboratory of Photobiology and Molecular Diagnostics, Intercollegiate Faculty of Biotechnology, University of Gdansk and Medical University of Gdansk, Gdańsk, Poland; Institute of Molecular Virology, University Hospital Ulm, Ulm, Germany; Department of Genomics, Medfuture Institute for Biomedical Research, Iuliu Hatieganu University of Medicine and Pharmacy, 400337 Cluj-Napoca, Romania; Institute of Biochemistry and Molecular Biology, Ulm University, Ulm, Germany; Department of Microbiology, Central University of Haryana, Mahendergarh, India; Central Facility for Electron Microscopy, Ulm University, Ulm, Germany

**Keywords:** Angicin, mannose phosphotransferase system, bacteriocin receptor, structure activity relationship

## Abstract

Angicin is a class IId bacteriocin produced by *Streptococcus anginosus* with activity against Gram-positive pathogens, including *Listeria monocytogenes* and vancomycin-resistant *Enterococcus faecium*. While the mannose phosphotransferase system (Man-PTS) has been identified as a receptor in *L. monocytogenes*, its role in streptococci and the structural determinants of Angicin activity remain unclear. Here, we demonstrate that the Man-PTS is required for Angicin susceptibility in *Streptococcus constellatus*. A transposon mutant (*manM*::ISS1) showed complete resistance to Angicin and impaired mannose utilization. Structure–activity relationship analysis of truncated and modified peptides localized antimicrobial activity to the C-terminal region, although none of the variants matched the activity of the full-length peptide. Angicin induced membrane depolarization and pore formation in target bacteria. Residual activity in Man-PTS-impaired *L. monocytogenes* suggests an additional receptor-independent effect at higher concentrations. *In vivo* toxicity analysis using zebrafish embryos showed low toxicity at active concentrations. These findings identify the Man-PTS as a receptor for Angicin in streptococci and define structural features associated with its antimicrobial activity.

## Introduction

The global emergence of antibiotic-resistant bacteria, accompanied by a rise in infection-related mortality, underscores the urgent need for novel antimicrobial therapeutics (*WHO Bacterial Priority Pathogens List 2024*, 2024). In 2019, antibiotic-resistant infections were responsible for 1.27 million deaths and contributed to 4.95 million deaths worldwide (Murray et al., 2022), with projections estimating up to 8.22 million deaths annually by 2050 (Kariuki, 2024). Consequently, the identification of safe, effective, and readily producible antimicrobial compounds represents a major priority. Bacteriocins, ribosomally synthesized antimicrobial peptides produced by bacteria, have emerged as promising candidates for clinical applications and food preservation (Darbandi et al., 2022; Mathur et al., 2017).

*Streptococcus anginosus*, an opportunistic pathogen commonly found in the oral cavity, produces the bacteriocin Angicin (Vogel et al., 2021). Angicin belongs to class IId bacteriocins, which are typically small (<10 kDa), structurally stable peptides with minimal post-translational modifications. Similar to other members of this class, Angicin has been shown to interact with the mannose phosphotransferase system (Man-PTS), which serves as a receptor in *Listeria monocytogenes* (Tymoszewska et al., 2017; Tymoszewska and Aleksandrzak-Piekarczyk, 2024; Vogel et al., 202*2)*. The Man-PTS is a phosphoenolpyruvate-dependent transport system responsible for the uptake and phosphorylation of carbohydrates such as glucose and mannose (Postma et al., 1993). In the context of bacteriocin activity, the N-terminal region of class IId bacteriocins mediates binding to extracellular loops of the Man-PTS, while the C-terminal α-helical region inserts into the transporter, keeping it in an open state and thereby causing membrane disruption (Li et al., 2023).

The Man-PTS is widely distributed among bacteria, including enterobacteriaceae (e.g. *Escherichia coli, Klebsiella pneumoniae*), lactic acid bacteria (e.g. streptococci, enterococci), and other bacillales such as staphylococci and listeria. In contrast, many non-fermenting pathogens (e.g. *Pseudomonas aeruginosa, Acinetobacter baumannii*) lack a functional Man-PTS. In *L. monocytogenes*, the system consists of four genes (*manA, manB, manC, and manD*), encoding cytoplasmic and membrane-associated subunits required for sugar transport (Postma et al., 1993; Stoll and Goebel, 2010). Mutations in the membrane-associated components ManC and ManD, particularly within the extracellular γ-loop region, are associated with decreased susceptibility to class IId bacteriocins (Oftedal et al., 2021; Tymoszewska et al., 2017; Vogel et al., 2022). In streptococci, homologous components are encoded by *manL* (cytoplasmic), *manM*, and *manN* (membrane-associated) (Asanuma et al., 2004; Sundar et al., 2017). While it is plausible that the Man-PTS also functions as a receptor for class IId bacteriocins in streptococci, this has not been experimentally validated.

Angicin exhibits strong antimicrobial activity against *L. monocytogenes*, vancomycin-resistant *Enterococcus faecium* (VRE), and several streptococcal species, whereas most ESKAPE pathogens (*Enterococcus faecium, Staphylococcus aureus, K. pneumoniae, A. baumannii, P. aeruginosa*, and *Enterobacter spp*.) are only affected at higher concentrations (Miller and Arias, 2024; Vogel et al., 2021). To enhance antimicrobial potency or broaden the spectrum of activity, bacteriocins have been subjected to structure–activity relationship (SAR) analyses and bioengineering approaches. Such strategies have yielded derivatives with improved efficacy, including modified variants of other bacteriocins, like nisin and mutacin (Guo et al., 2024; Kers et al., 2018). However, for Angicin, the structural determinants underlying its antimicrobial activity have not been systematically investigated.

Taken together, it remains unclear whether the Man-PTS mediates Angicin susceptibility in streptococci and which structural features of Angicin are required for its antimicrobial activity. In this study, we therefore investigated the molecular basis of Angicin function. We generated a transposon mutant library in *Streptococcus constellatus* to identify factors required for susceptibility and performed a structure–activity relationship analysis of Angicin using truncated and modified peptide variants. In addition, we analyzed membrane effects and evaluated *in vivo* toxicity using a zebrafish embryo model.

## Results

### Angicin induces membrane depolarization and pore formation

Angicin is the first and thus far the only bacteriocin described in *S. anginosus*. Based on previous experiments using SYTOX, liposomes and pHluorin assays, it was shown to disrupt the membrane of bacterial target cells. To investigate whether this damage also includes membrane depolarization, we performed a DiBAC_4_(3) assay. In these experiments, Angicin concentrations as low as 6.25 µg/ml for VRE (*p* = 0.0152) and 0.2 µg/ml for *L. monocytogenes* (*p* = 0.0303) led to a significant membrane depolarization within 30 min (Fig. 1a). However, the overall effect on membrane potential remained relatively moderate, and even at 100 µg/ml, depolarization did not reach the levels observed for the positive control (heat-treated bacteria; *p* = 0.0043).

**Figure 1:**
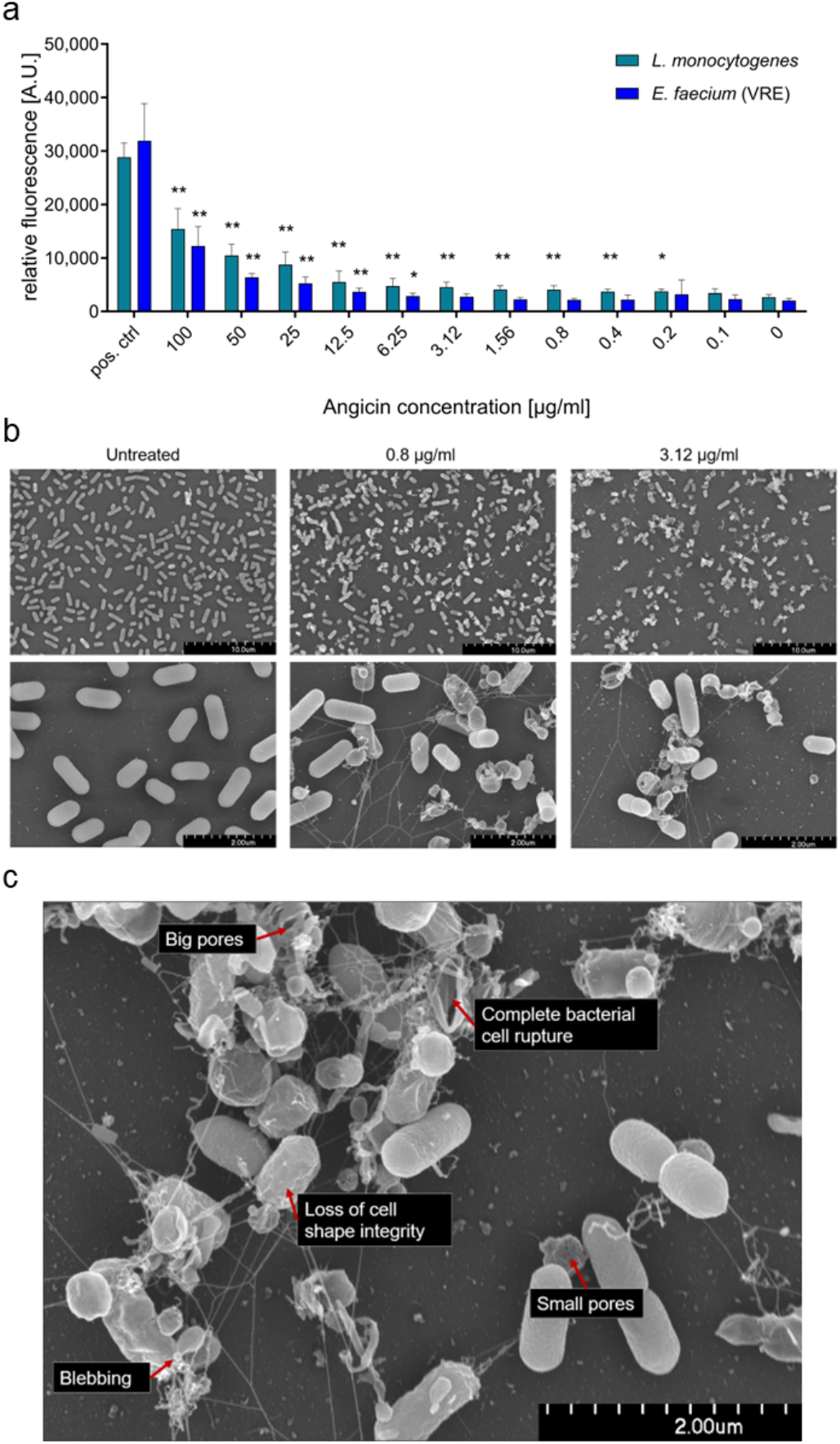
Angicin leads to pore formation in the membrane of target cells. a) *L. monocytogenes* or *E. faecium* (VRE) were incubated with Angicin for 30 min. Subsequently, a DiBAC_4_(3) –assay was used to assess effects on membrane polarization. As a positive control heat-treated bacteria were used. Depicted is the mean + standard deviation (S.D.) of five independent experiments. To calculate significant differences to untreated bacteria, a Mann-Whitney U test was used (* p-value < 0.05, ** p-value < 0.01). *L. monocytogenes* was incubated with 3.125, 0.8 or 0 µg/ml of Angicin for 24 h; afterwards bacterial cells were fixed and analysed by scanning electron microscopy. At least 14 pictures per treatment condition were taken. b) Representative pictures are depicted to give an overview over the whole sample. c) *L. monocytogenes* was treated with 3.125 µg/ml Angicin for 24 h. The different stages of pore formation are illustrated and labelled.

To assess whether structural membrane damage occurs at concentrations inducing depolarization, scanning electron microscopy (SEM) was performed. Treatment of *L. monocytogenes* with 0.8 µg/ml or 3.12 µg/ml Angicin for 24 h resulted in pronounced morphological alterations, including loss of cell shape integrity, membrane blebbing, leakage of intracellular content, and formation of membrane pores (Fig. 1b,c). In some cases, complete rupture of bacterial cells was observed, whereas untreated controls remained intact.

Together, these results demonstrate that Angicin induces membrane depolarization and visible membrane damage at low concentrations, consistent with pore formation in bacterial membranes. Given that class II bacteriocins frequently rely on specific membrane receptors such as the Man-PTS to initiate pore formation, we next investigated whether Angicin activity depends on this system.

### Man-PTS is required for Angicin-mediated killing of *Streptococcus constellatus*

Antimicrobial activity against *L. monocytogenes* relies on the interaction of Angicin with the Man-PTS (Vogel et al., 2022). While Man-PTS as a class IId bacteriocin receptor has been extensively studied in *L. monocytogenes*, it has rarely been investigated in other species such as streptococci (Tymoszewska and Aleksandrzak-Piekarczyk, 2024). Since Angicin shows potent activity against *S. constellatus*, we aimed to identify its receptor in this species (Fig. 2a).

**Figure 2:**
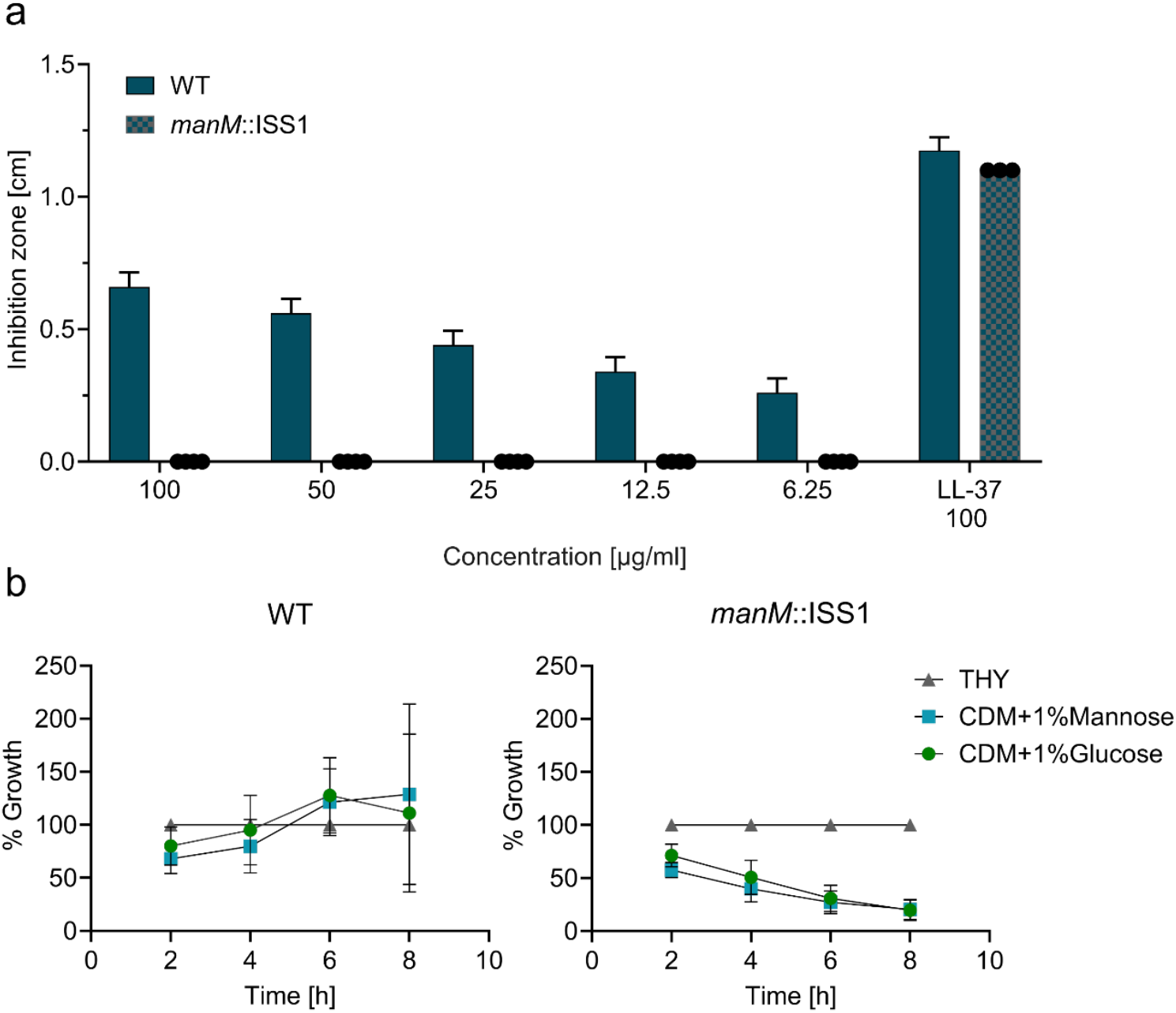
Analysis of *S. constellatus* SK53 Man-PTS mutants on (a) sensitivity towards Angicin and (b) on sugar metabolism. a) Wild type (WT) and mutants were tested in a radial diffusion assay for susceptibility towards Angicin in five independent experiments. LL-37 was used as a positive control. b) To assess the functionality of the Man-PTS, WT and *manM*::ISS1 were grown in chemically defined medium with or without 1% D-mannose or D-glucose. Growth was monitored over a period of 8 h spectrometrically. For comparison, the strains were additionally incubated in THY. Growth was normalized to growth in THY. At least five independent experiments were performed.

By generating a mutant library in the Angicin-sensitive *S. constellatus* wild type strain SK53 and selecting for resistant isolates at 100 µg/ml Angicin, a resistant clone was identified. Mapping of the insertion site revealed disruption at position 586 bp within the *manM* gene (806 bp), likely exerting a polar effect that disrupts expression of the entire operon. Following plasmid clearance, the resulting SK53*manM*::ISS1 mutant was tested for sensitivity towards the Angicin-producing strain *S. anginosus* BSU 1211 and synthetic Angicin in radial diffusion assays (RDA).In both assays, the Man-PTS mutation led to a complete loss of susceptibility towards Angicin (Supplementary Table 1, Fig. 2a), demonstrating that an intact Man-PTS is required for Angicin activity against *S. constellatus*.

**Table 1:**
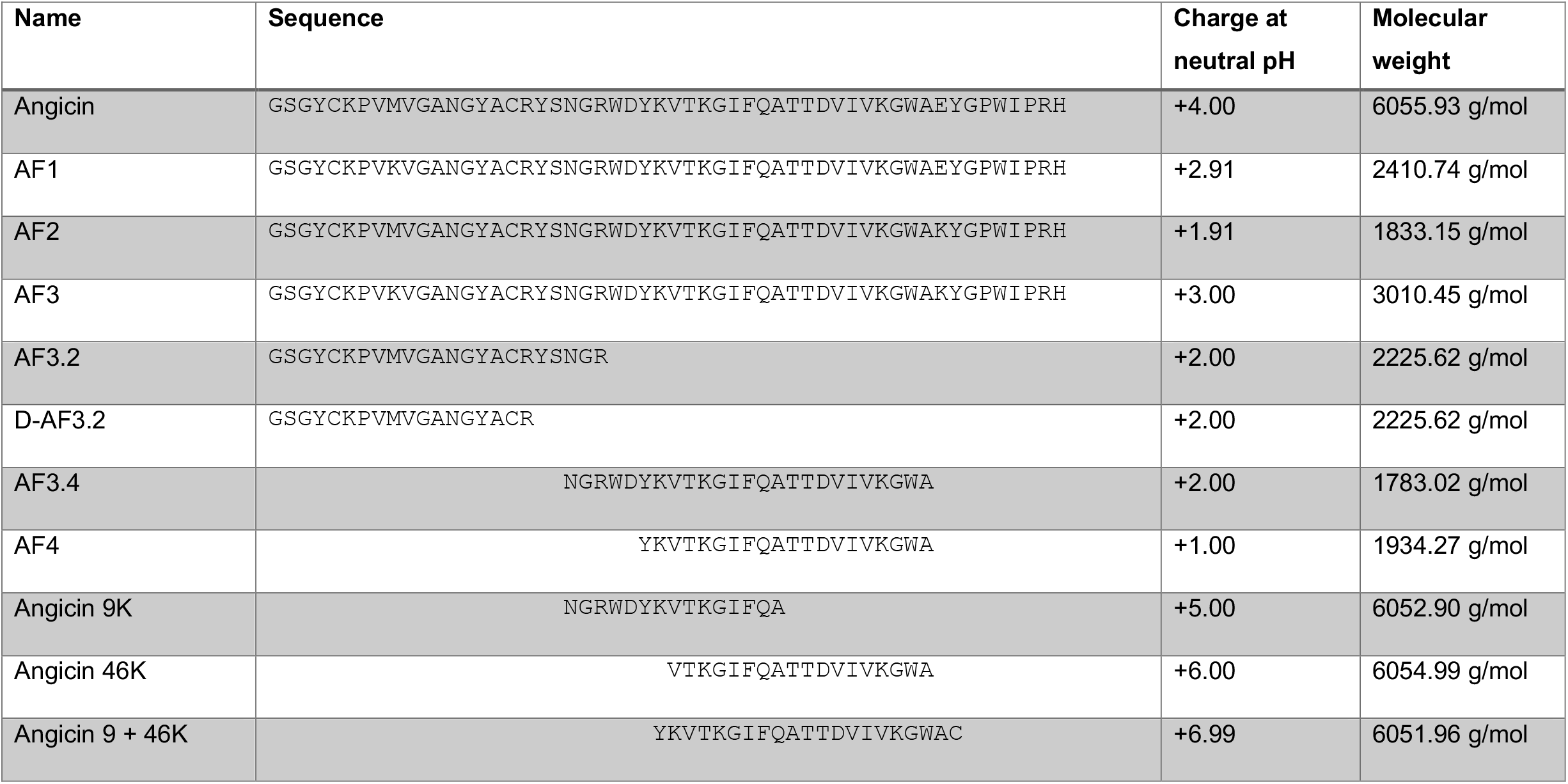
Sequence, charge and molecular weight of Angicin derivatives used in this study. Molecular weight and net charge were calculated using Bachem calculator. Bachem peptide calculator accessed on 21.06.2024 at 9.03 am. Angicin fragments are abbreviated as AF.

In a second step, the transporter subunits of Angicin-sensitive and -insensitive species were compared with the Man-PTS of *S. constellatus* (Supplementary Table 2). As expected, subunits IIC and IID of *S. constellatus* and *S. anginosus* showed high similarity (87.31% and 86.47%, respectively). For *L. monocytogenes* and *E. faecium*, sequence identities of 75.37% and 73.61% were observed. In contrast, *E. coli* and *K. pneumoniae* exhibited sequence identities below 50%, and both species possess an IID subunit lacking a γ-loop.

Further investigations verified the successful disruption of the Man-PTS gene locus in *S. constellatus* by assessing its physiological consequences. Wild type and the *manM* mutant strain were cultivated in chemically defined medium (CDM, Supplementary Method) supplemented with different carbon sources. While neither strain was able to grow in the absence of sugar, the wild type exhibited robust growth in the presence of mannose and glucose (Fig. 2b). In contrast, the SK53*manM*::ISS1 mutant showed significantly impaired growth, reflected by markedly reduced OD_600_ values after 6 h (D-mannose: *p* = 0.0079; D-glucose: p = 0.0079) and 8 h (D-mannose: *p* = 0.0411; D-glucose: p = 0.0649, not significant). These results demonstrate that disruption of the *manM* locus resulted in a clear growth defect under carbohydrate-dependent conditions.

In conclusion, these results demonstrate that the Man-PTS transporter is essential for Angicin susceptibility in *S. constellatus*, identifying it as a critical determinant of Angicin-mediated killing. These findings raise the question which structural elements of Angicin mediate receptor interaction versus membrane disruption, and whether these functions can be uncoupled.

### Structure–activity relationship (SAR) analysis of Angicin

To delineate the structural determinants underlying receptor interaction and membrane disruption, we performed a systematic structure–activity relationship (SAR) analysis of Angicin. For this, truncated fragments and modified variants were generated and tested for antimicrobial activity against established Angicin target species as well as ESKAPE pathogens using radial diffusion assays.

Potentially active truncated variants of Angicin were predicted using CAMP4_R4_, ClassAMP, and AMPscanner (Fig. 3, Table 1), synthesized, and experimentally evaluated. To assess how these modifications affect peptide structures, AlphaFold modelling was performed (Fig. 4a). The predicted structure of full-length Angicin consists of three antiparallel β-sheets at the N-terminus and an α-helical C-terminus. Accordingly, Angicin fragments (AF) 1 and 2, covering the N-terminal region, adopt β-sheet structures, whereas AF3 and AF4, derived from the C-terminal region, are predicted to form α-helices (Fig. 4a).

**Figure 3:**
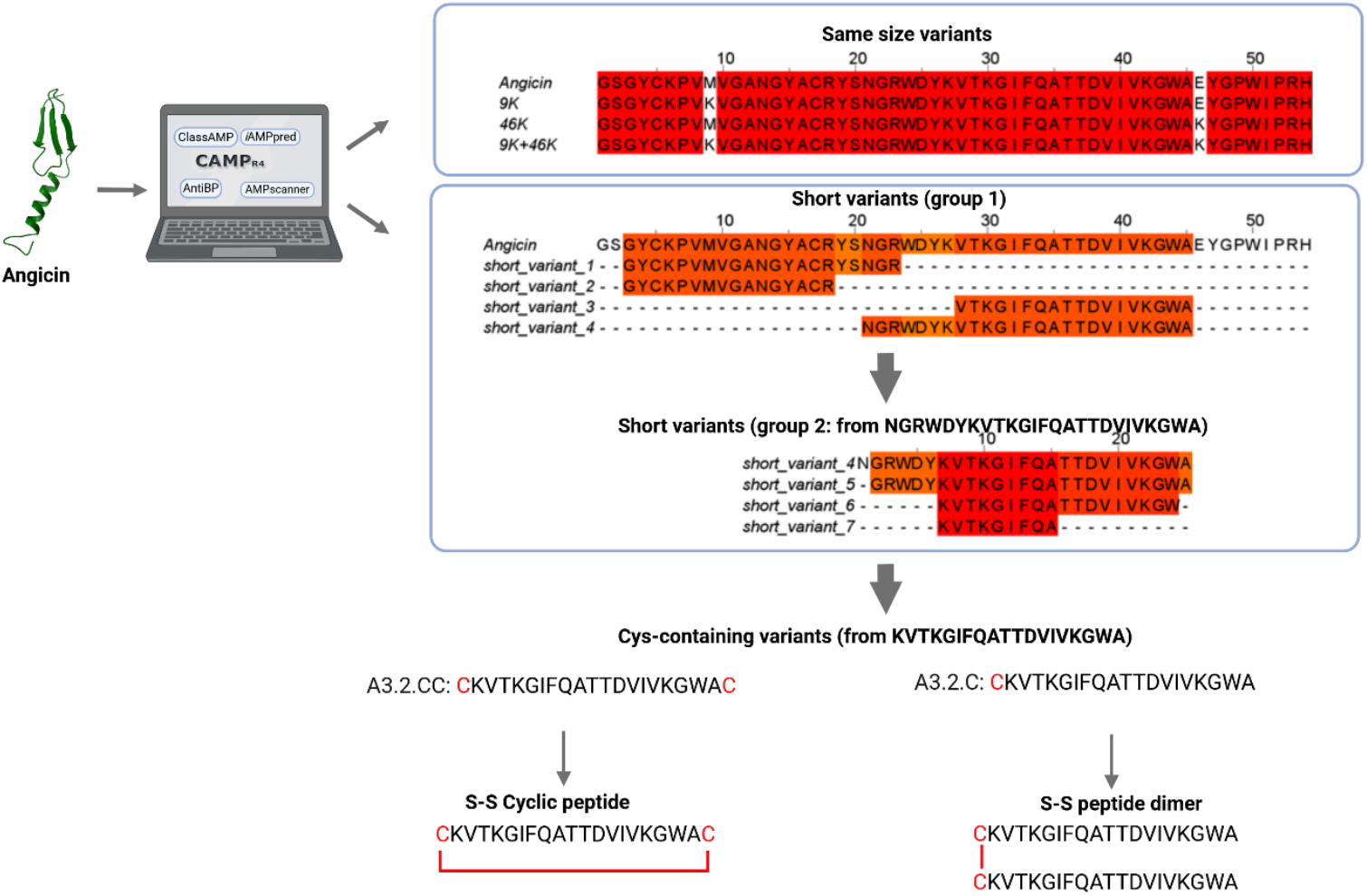
Overview of bioinformatic strategy and predicted peptides. Starting from the native Angicin sequence, antimicrobial prediction algorithms (CAMP_R4_, ClassAMP, and AMPscanner) guided the design of: (1) same-size variants with single (46K) or double (9K+46K) amino acid substitutions, (2) truncated peptides derived from the N-terminal region (group 1) or the C-terminal bioactive core NGRWDYKVTKGIFQATTDVIVKGWA (group 2), and (3) cysteine-containing variants for cyclization (A3.2.CC) or dimerization (A3.2.C).

**Figure 4:**
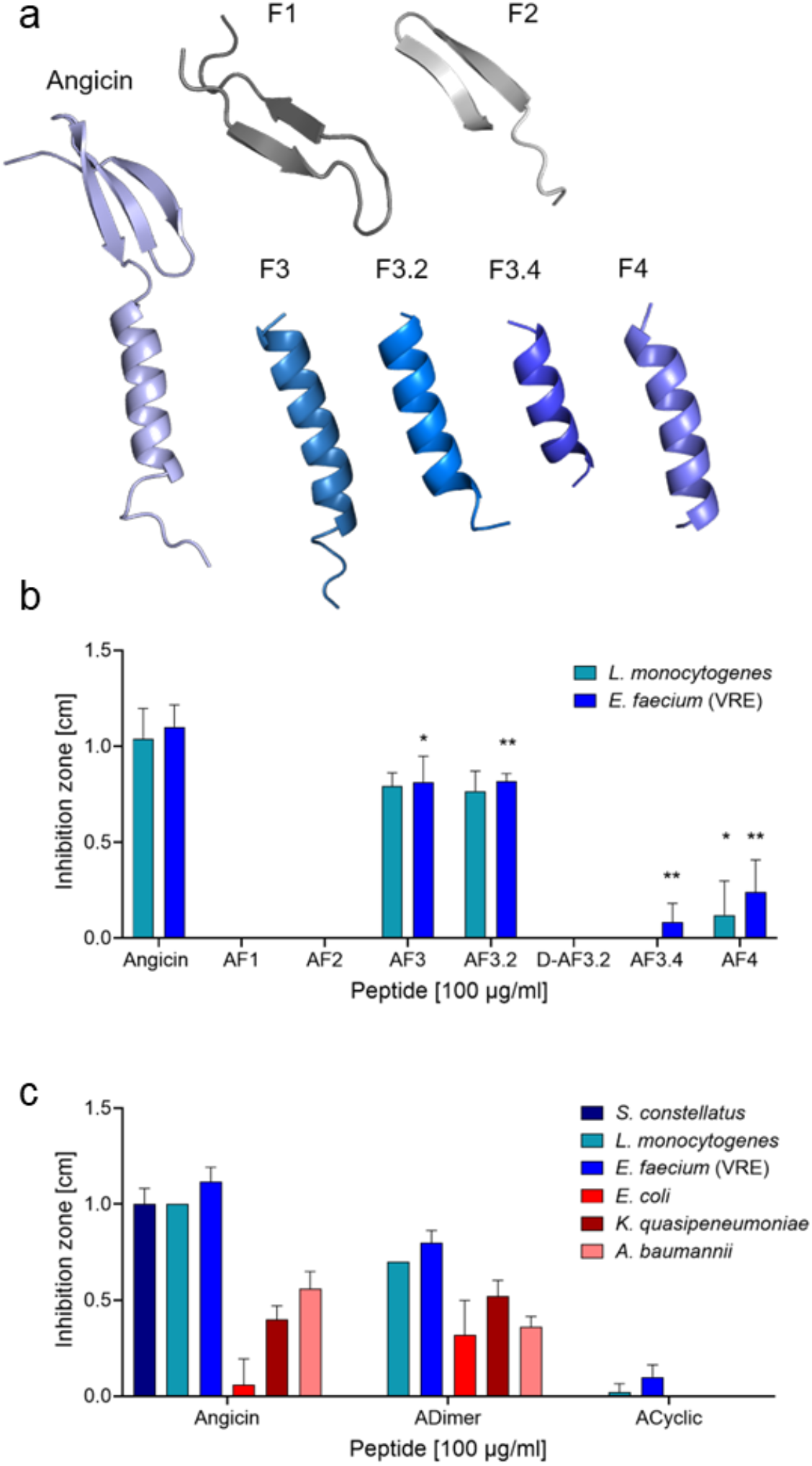
Characterization of short Angicin variants. a) Alphafold structure of all variants was predicted with ColabFold 2.0. b) The spectrum of activity of linear Angicin fragments was determined using a radial diffusion assay against *L. monocytogenes* and VRE. c) The spectrum of activity of structurally different Angicin variants. b+c: Depicted is the mean + S.D. of at least five independent experiments. Significant differences were calculated with a Mann-Whitney U test (* indicates p-value < 0.5, ** illustrates p-value < 0.1).

Functional analysis revealed that AF1 and AF2 did not exhibit antimicrobial activity, whereas AF3 showed the strongest inhibitory effect against *L. monocytogenes* and vancomycin-resistant *E. faecium* (VRE). AF4 displayed only minimal activity (Fig. 4b). Further truncation of AF3 identified AF3.2 as the shortest fragment retaining measurable activity, while AF3.4 was inactive. However, AF3.2 displayed reduced potency compared to full-length Angicin and a restricted activity spectrum, with inhibition observed for *L. monocytogenes* and VRE, and only weak activity against *P. aeruginosa*. No activity was detected against *E. coli, S. aureus, K. pneumoniae, A. baumannii*, or streptococcal strains (Supplementary Table 1). To further explore structural requirements, AF3.2 variants were modified by incorporation of D-amino acids, cyclization, or dimerization. Incorporation of D-amino acids resulted in a complete loss of activity, while cyclization retained only minimal activity (Fig. 4b,c). In contrast, dimerization altered the activity profile, resulting in reduced activity against Gram-positive bacteria but increased activity against Gram-negative species, including *E. coli, A. baumannii, K. quasipneumoniae*, and *P. aeruginosa*.

### Antimicrobial activity of modified full-length Angicin variants

Since truncated variants did not reach the activity of full-length Angicin, further optimization focused on targeted amino acid substitutions within the full-length peptide. Based on structural considerations, methionine at position 9 and glutamate at position 46 were replaced by lysine, generating the variants Angicin 9K, 46K, and 9+46K (Fig. 3; Table 1). These substitutions increased the net positive charge of the peptide (Table 1). While all variants retained comparable activity against Gram-positive bacteria, no improvement over wild-type Angicin was observed (Fig. 5a). However, an enhanced activity against Gram-negative bacteria, including *E. coli* and *K. quasipneumoniae*, was observed for variants 46K (p = 0.0238; p = 0.0159) and 9+46K (p = 0.0238; p = 0.0317). This was supported by liposome leakage assays using liposomes formed from lipids derived from *E. coli* and *P. aeruginosa* cultures, where these variants showed increased membrane-disrupting activity compared to wild-type Angicin (Fig. 5b).

**Figure 5:**
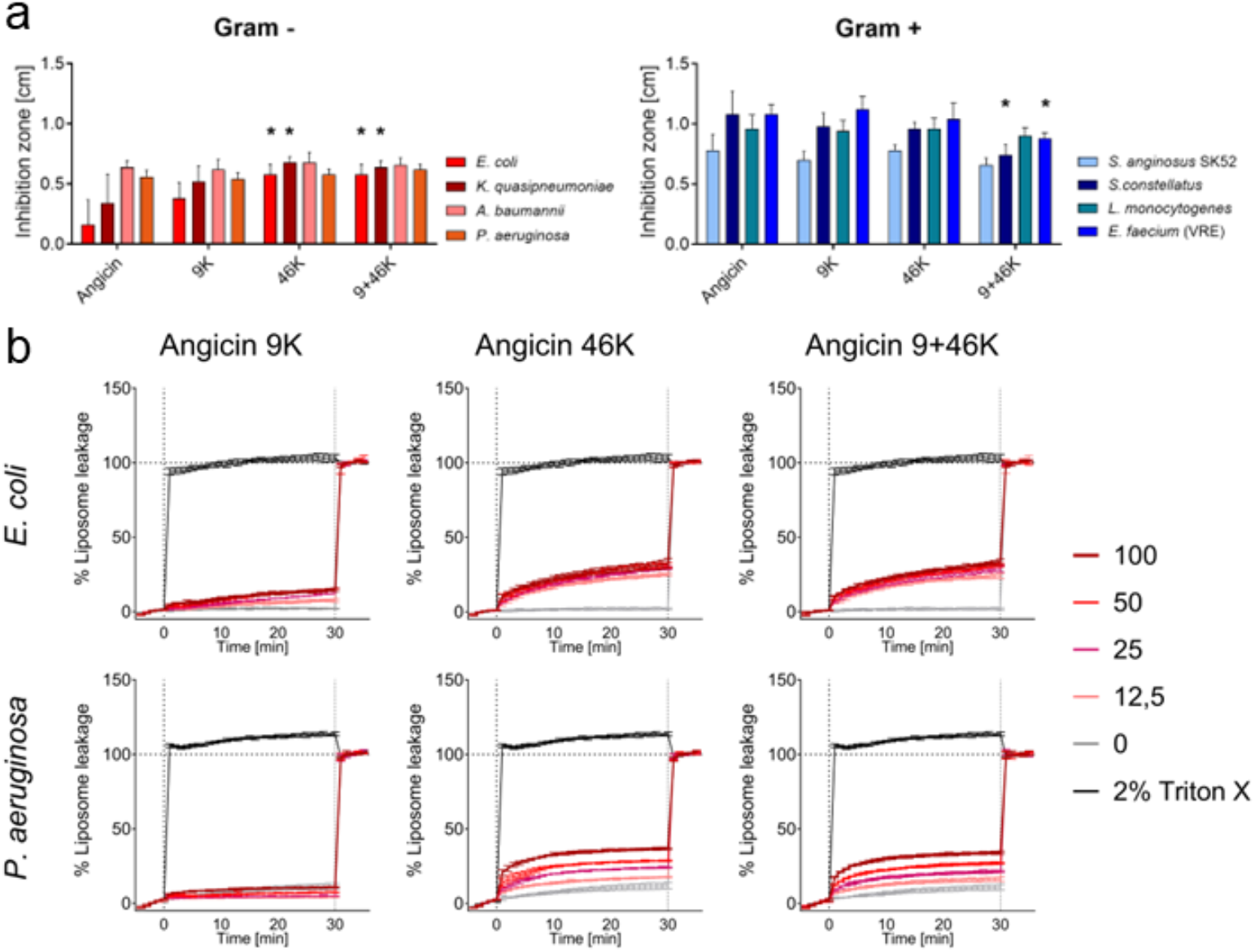
Characterization of Angicin variants. a) The antimicrobial activity of Angicin variants was tested in a radial diffusion assay against *S. anginosus, S. constellatus, L. monocytogenes* and the ESKAPE pathogens. Illustrated is the mean + S.D. of at least five independent experiments. Significant differences to Angicin were calculated with a Mann-Whitney U test (* p-value <0.05; ** p-value < 0.01). b) A liposome leakage assay was performed with *E. coli* and *P. aeruginosa* derived liposomes, which were treated with the Angicin variants. Depicted is the mean ± S.D. of three experiments.

Collectivley, these results demonstrate that the C-terminal region of Angicin is critical for antimicrobial activity, while the N-terminal region alone is insufficient for bacterial killing. Moreover, both sequence composition and structural integrity are required for full activity, as none of the truncated or modified variants matched the potency of the full-length peptide. Notably, the observation that truncated variants retain partial membrane activity but lose full antimicrobial potency suggests that Angicin function involves both receptor-dependent and receptor-independent mechanisms.

### Angicin shows a favorable *in vivo* toxicity profile in zebrafish embryos

Given the identification of active Angicin variants and the importance of preserving activity while minimizing cytotoxicity, we next assessed the *in vivo* toxicity profile of Angicin using a zebrafish embryo model. Zebrafish embryos (*Danio rerio*) were exposed to Angicin at concentrations ranging from 0.3 to 30 µM (1.8–181 µg/ml) for 24 h and subsequently analyzed for toxic effects (Fig. 6, Supplementary Fig. 1). Due to their optical transparency, zebrafish embryos enable the simultaneous assessment of multiple toxicity parameters, including general cytotoxicity (necrosis, lysis), developmental toxicity (developmental delay, malformations), cardiotoxicity (e.g., edema, impaired circulation), and neurotoxicity (reduced touch response) (Supplementary Fig. 1) (Morash et al., 2011a; Noschka et al., 2021). At concentrations up to 3 µM, no signs of toxicity were observed (Fig. 6). At the highest concentration tested (30 µM), mild cardiotoxicity and general cytotoxic effects were detected. Importantly, no neurotoxicity or developmental toxicity was observed at any concentration. Together, these results demonstrate that Angicin exhibits low *in vivo* toxicity within the concentration range associated with antimicrobial activity, supporting its potential for further development.

**Figure 6:**
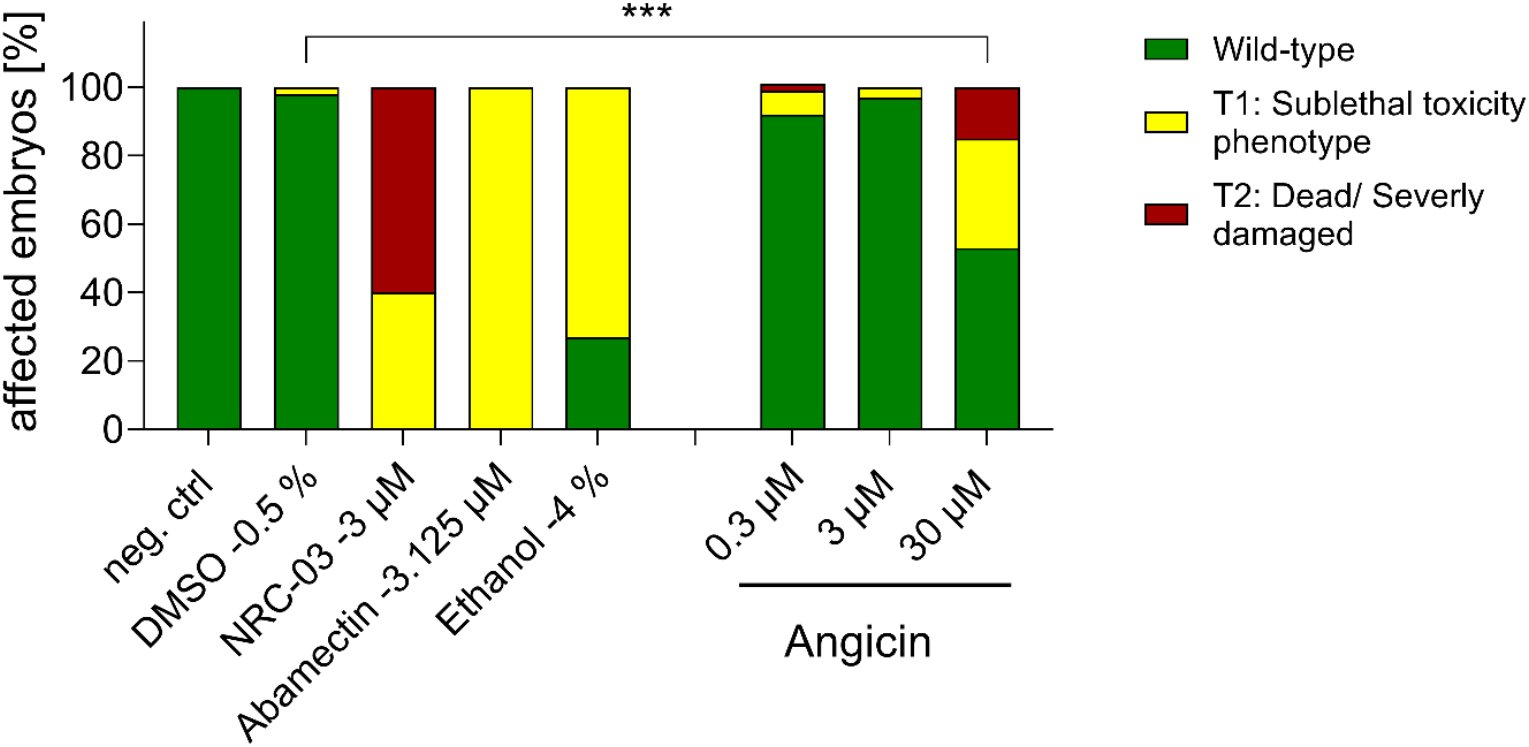
Toxicity of Angicin in zebrafish embryos. Zebrafish embryos (*Danio rerio*) were exposed to Angicin (0.3-30 µM) for 24h. Subsequently, cytotoxicity, cardiotoxicity, developmental and neurotoxicity were analysed with a stereomicroscope and touch response. Depicted is a summary of the overall toxicity. In total, 60 embryos were exposed to each Angicin concentration in two independent experiments. As neg. ctrl, embryo medium was used. NRC-03, Abamectin and Ethanol serve as positive controls for cytotoxicity, neurotoxicity and cardiotoxicity, respectively. A Chi-square test was used to investigate significant differences, *** indicates a p-value below 0.001.

### Differential dependence of Angicin activity on the Man-PTS across bacterial species

To directly test whether Angicin activity differentially depends on the Man-PTS across bacterial species, *L. monocytogenes* Δ*manD* and *S. constellatus SK53manM::ISS1* mutants, along with their respective wild-type strains, were exposed to full-length Angicin and Angicin fragments (Fig. 7). It should to be noted, that while we have a complete disruption of the Man-PTS in *S. constellatus SK53manM::ISS1*, in *L. monocytogenes* Δ*manD* only 28 amino acids of the γ-loop are deleted.

**Figure 7:**
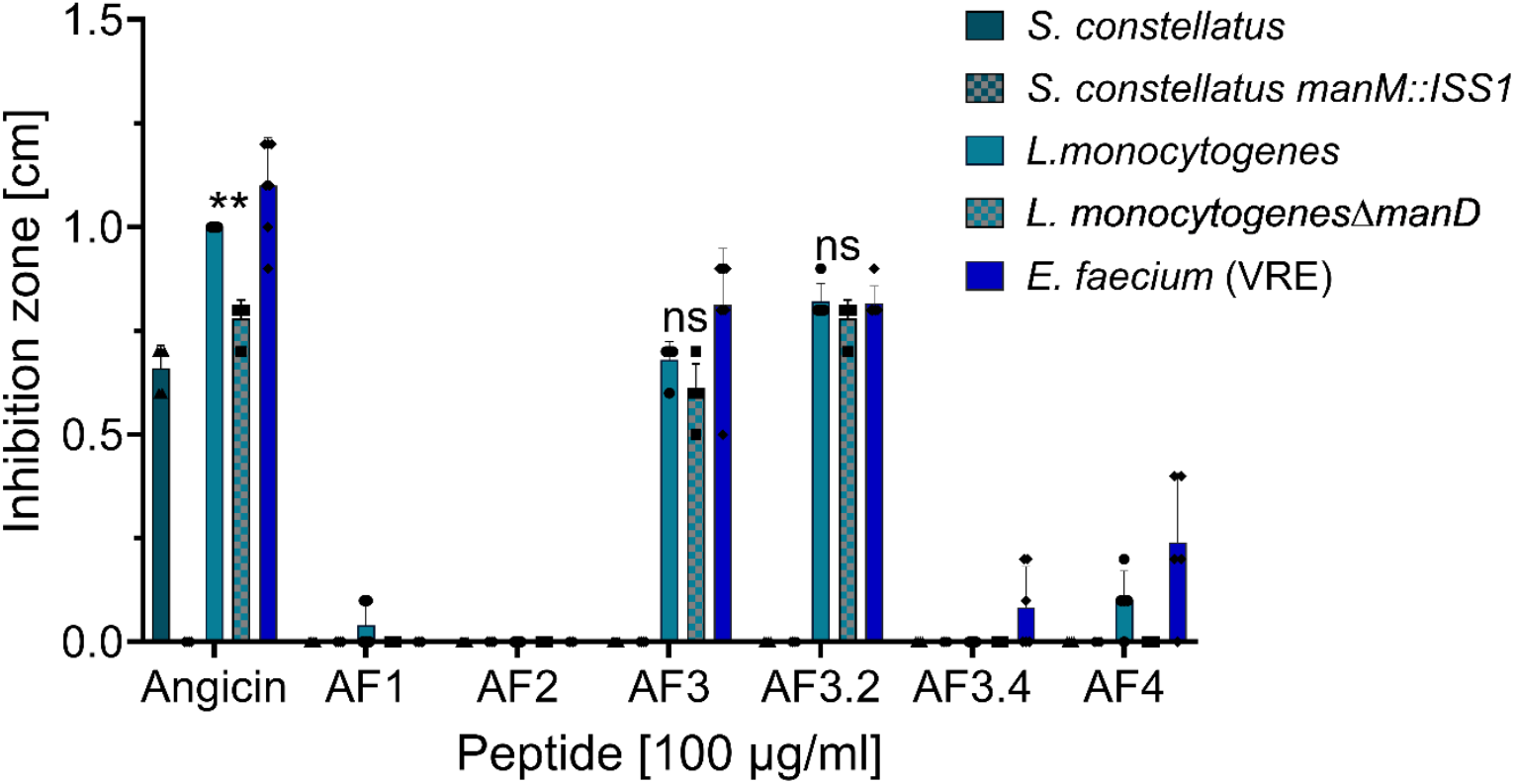
Influence of the Man-PTS on Angicin fragment sensitivity. *L. monocytogenes* and S. *constellatus* and their respective Man-PTS mutants were tested in a radial diffusion assay for sensitivity towards Angicin fragments. Depicted is the mean + S.D. of at least five experiments. Inactive peptides were tested at least two times. Significant differences were calculated using a Mann-Whitney U-test (p-value <0.05 is depicted by *; p-value < 0.01 is illustrated with **).

In *L. monocytogenes*, Angicin fragments displayed comparable inhibitory activity against both the wild-type and the *ΔmanD* mutant strain, although overall activity was reduced compared to full-length Angicin in the wild type. Importantly, the wild-type strain remained significantly more susceptible to full-length Angicin than the *ΔmanD* mutant (*p* = 0.0079), confirming that the Man-PTS acts as a major receptor enhancing Angicin activity, but is not strictly required for membrane disruption.

In contrast, in *S. constellatus*, full-length Angicin exhibited strong antimicrobial activity against the wild-type strain, whereas the Man-PTS mutant was completely resistant (Fig. 7). Notably, neither the wild type nor the mutant strain showed any sensitivity to Angicin fragments. Collectively, these results indicate that Angicin activity in *S. constellatus* is strongly associated with an intact Man-PTS. In *L. monocytogenes*, the residual activity observed in the Δ*manD* mutant and following treatment with Angicin fragments suggests that, in addition to Man-PTS-dependent interactions, Angicin may exert antibacterial effects through an additional mechanism contributing to membrane damage.

## Discussion

Angicin is a type IId bacteriocin produced by *S. anginosus*, targeting Gram-positive bacterial species while displaying high structural stability and low cytotoxic (Vogel et al., 2022, 2021). As previously reported for Angicin and other class IId bacteriocins, its antimicrobial activity relies on disruption of membrane integrity of target bacteria (Héchard and Sahl, 2002; Tymoszewska et al., 2017; Vogel et al., 2022, 2021). The aim of this study was to further characterize the molecular determinants underlying Angicin-mediated membrane disruption and receptor interaction.

Building on previous work demonstrating membrane damage (Vogel et al., 2022, 2021), we show that Angicin induces membrane depolarization and structural membrane disruption, as visualized by SEM. The observed morphological changes—including loss of cell integrity, membrane blebbing, and pore formation—represent characteristic antimicrobial peptide (AMP)-mediated pore formation (Deshmukh et al., 2021; Hartmann et al., 2010). These findings are consistent with established models of AMP-induced membrane disruption, including both barrel-stave and toroidal pore formation mechanisms (Mahlapuu et al., 2016). However, the relatively moderate depolarization observed even at high Angicin concentrations suggests that membrane disruption is not purely detergent-like but instead involves the formation of defined pore structures.

Type IId bacteriocins typically induce pore formation through interaction with the Man-PTS, which acts as a membrane receptor (Tymoszewska and Aleksandrzak-Piekarczyk, 2024). In line with this, our data demonstrate that Angicin activity in *S. constellatus* strictly depends on an intact Man-PTS. Disruption of the Man-PTS operon resulted in complete resistance, confirming that Man-PTS functions as an essential receptor in streptococci. While Man-PTS-mediated bacteriocin activity has been extensively characterized in *L. monocytogenes* (Tymoszewska et al., 2017; Tymoszewska and Aleksandrzak-Piekarczyk, 2024; Vogel et al., 2022), its functional role in streptococci has remained largely unproven. Our findings therefore extend the role of Man-PTS as a conserved receptor for class IId bacteriocins across different bacterial genera.

At the molecular level, the Man-PTS is a phosphoenolpyruvate-dependent transport system responsible for carbohydrate uptake and phosphorylation (Postma et al., 1993). In the context of bacteriocin activity, it has been proposed that the N-terminal region of class IId bacteriocins mediates binding to the extracellular γ-loop of the Man-PTS, while the C-terminal α-helical region inserts into the membrane and induces pore formation (Li et al., 2023). Our SAR data support this model. Truncation experiments revealed that the C-terminal region of Angicin is critical for antimicrobial activity, whereas the N-terminal region alone is insufficient to mediate bacterial killing. At the same time, truncated variants retained partial membrane-disruptive activity, indicating that membrane interaction and receptor engagement represent separable functional components.

Interestingly, Angicin retained substantial antimicrobial activity in *L. monocytogenes* strains carrying mutations that impair Man-PTS function. This finding suggests that, unlike in streptococci, Angicin activity in *L. monocytogenes* may not be exclusively dependent on an intact Man-PTS receptor. Rather, Angicin appears capable of exerting antimicrobial effects through both receptor-mediated interactions and direct membrane perturbation. While further studies are required to elucidate the underlying mechanism, these results suggest a more flexible mode of action for Angicin than has previously been recognized for class IId bacteriocins. Such differences may arise from variations in membrane composition, receptor accessibility, or peptide–membrane interactions between bacterial species.

Further insights into structure–function relationships were obtained through engineering of Angicin variants. While truncated peptides did not reach the activity of the full-length peptide, charge-enhanced variants (e.g., 46K, 9+46K) showed increased activity against Gram-negative bacteria, including *E. coli* and *K. quasipneumoniae*. This is consistent with the known role of electrostatic interactions in antimicrobial peptide activity, where increased cationicity promotes interaction with negatively charged bacterial membranes (Vejzovic et al., 2022).

Importantly, the *in vivo* toxicity assessment using zebrafish embryos confirmed that Angicin exhibits low toxicity within the therapeutically relevant concentration range of 0.3-3 µM. Only at the highest concentrations tested were mild cardiotoxic and general cytotoxic effects observed, while neurotoxicity and developmental toxicity were absent. These findings support the notion that receptor-mediated targeting contributes to selectivity and suggest that Angicin maintains a favorable therapeutic window.

Taken together, our findings support a model in which Angicin combines Man-PTS–mediated targeting with C-terminal membrane pore formation, with the relative contribution of these mechanisms depending on the bacterial species. In streptococci, Angicin activity is strictly receptor-dependent, whereas in *L. monocytogenes*, receptor-independent membrane interactions contribute to residual activity. This dual-mode mechanism provides both specificity and robustness and may reduce susceptibility to resistance development.

From a broader perspective, this work refines the current understanding of class IId bacteriocins by demonstrating that receptor dependence is not absolute but can be complemented by intrinsic membrane activity. The identification of functionally distinct regions within Angicin provides a framework for rational engineering of bacteriocins with tailored activity spectra and improved therapeutic properties.

## Material and Methods

Bacterial strains and peptides used in this study are listed in Supplementary Table 3 and Table 1, respectively.

### Bacterial culturing

Bacterial cultivation was carried out on Tryptone soya agar supplemented with 5 % sheep blood (Oxoid) at 37 C and 5% CO_2_. For liquid cultivation, *E. coli, P. aeruginosa, A. baumannii* were inoculated in lysogeny broth (LB-Miller) and incubated aerobically at 37 °C with shaking (160 rpm). Listeria strains were either grown in brain-heart-infusion (Oxoid) or Todd-Hewitt broth (Oxoid) with 0.5 % yeast (THY, Gibco). For the liquid cultivation of streptococci, enterococci, staphylococci and klebsiella, THY was used as growth medium, and incubation was performed at 37 °C and 5 % CO_2_.

### DiBAC_4_(3) -Assay

Overnight cultures of *L. monocytogenes* and VRE were used to inoculate fresh cultures in the morning with an O.D. _600 nm_ of 0.02. After bacterial cells reached mid-exponential phase (O.D. _600 nm_ above 0.3) the equivalent of an O.D. _600 nm_ 0.3 in 350 µl was transferred to a reaction tube and centrifuged at 17 000 x g for 2 min. The pellet was resuspended in PBS with 5 µM Bis-(1,3-Dibutylbarbituric Acid) Trimethine Oxonol (DiBAC_4_(3), Invitrogen) and Angicin in concentrations ranging from 100 to 0.2 µg/ml. Bacterial cells without peptide were used as a negative control. Afterwards, incubation at 37 °C for 30 min followed. As a positive control bacteria were incubated at 70 °C instead of 37 °C. Assays were kept in the dark. After incubation, centrifugation at 4500 x g for 10 min followed. The pellet was washed with PBS and subsequently centrifuged under the same conditions. The pellet was resuspended in 350 µL PBS, and fluorescence was measured at 493 nm excitation and 525 nm emission using 100 µL aliquots in triplicate on an Infinite M200 Pro microplate reader (Tecan Group Ltd.).

### Scanning electron microscopy

An overnight culture of *L. monocytogenes* was adjusted to an O.D. _600 nm_ of 0.01 in THY supplemented with Angicin (3.125, 0.78, or 0 µg/ml). 2 ml of the bacterial suspension was placed in a well of a 12-well-plate and incubated for 24 h at 37 °C and 5 % CO2. The following day, the medium was replaced with fixing solution (2.5 % Glutaraldehyde, 0.1 mM phosphate-cacodylatbuffer, 1 % saccharose, pH 7.3) and kept overnight at 4 °C. Further preparation was performed as described previously (Schütz et al., 2021). In detail, samples were post-fixed with 2 % osmiumtetroxide in PBS for 20 min, washed in PBS, gradually dehydrated with increasing isopropanol concentrations, critical-point dried using carbon dioxide. Afterwards, samples were coated with about 2 nm of platinum-carbon by rotary electron beam evaporation and imaged in an Hitachi S-5200 SEM using the secondary electron signal. At least 14 images per condition were taken.

### Mutant library

A mutant library in *S. constellatus* SK53 was constructed using the vector pGhost9:ISS1, a plasmid combining a thermosensitive replication in streptococci at 30 °C with a chromosomal integration of the complete plasmid at 37 °C (Asam et al., 2013; Maguin et al., 1996; Spellerberg et al., 1999). Transformation of pGhost9:ISS1 was performed via induction of natural competence with competence-stimulating peptide-1, as previously described for *S. anginosus* (Bauer et al., 2018). Mutant library construction was performed, as described by Spellerberg et al. 1999 (Spellerberg et al., 1999).

Subsequently, the library was screened for Angicin-resistant mutants by incubation in THY supplemented with 100 µg/ml Angicin for 2 h. Angicin-treated cells were spread on sheep blood agar plates. *S. constellatus* mutants growing on this plate were checked for Angicin sensitivity by RDA. Finally, the DNA of resistant mutants was isolated (GenElute™ Bacterial Genomic DNA Kit, Sigma Aldrich, following the manufacturer’s instructions) and screened for an ISS1 insertion site in the Man-PTS operon. The operon was amplified using the AllTaq Master Mix Kit (Quiagen) according to the manufacturer’s protocols. Primers are listed in Supplementary Table 4. PCR products were purified with NucleoSpin® Gel and PCR Clean Up Kit (Machery-Nagel) and sequenced. Sequencing was performed by Microsynth Seqlabs.

### Growth assay

Chemically defined medium (CDM) was adapted from a formulation originally developed for *S. pyogenes* (Supplementary Method; Chang et al., 2011; Van De Rijn and Kessler, 1980). Overnight cultures of *S. constellatus* or its isogenic Man-PTS mutant were inoculated with a final O.D. _600 nm_ of 0.02 in CDM supplemented with either 1 % D-mannose, 1% D-glucose or water (neg. control). As a positive control bacteria were inoculated in THY. Growth was assessed spectrometrically (UV spectrophotometer UV-1800, Shimadzu) at 2,4,6 and 8 h post inoculation.

### Radial diffusion assay

#### Overlay Assay

A two-layer radial diffusion assay was used to assess the antimicrobial activity of peptides. As previously described (Vogel et al., 2021), overnight cultures of putative target strains were adjusted and inoculated into liquid agarosea at a density of 2 x 10^7^ bacterial cells per plate. Wells were placed into the plate and filled with 10 µl of a peptide. After 3 h of incubation at 37 °C, an overlay with trypticase soy agar was conducted. Measurement of inhibition zones was performed after overnight incubation at 37 °C and 5% CO_2_. Due to the slow growth of *S. constellatus*, wild type and *manM* mutant were analysed after 3 days of incubation at 37 °C and 5% CO_2_. As a positive control, 100 µg/ml of the antimicrobial peptide LL-37 (Anaspec) or nisin (Sigma) were used (Bucki et al., 2010; Gharsallaoui et al., 2016). All peptides investigated by this method are listed in Table 1. Peptides were purchased form PSL Heidelberg or ULMTec Core Facility Functional Peptidomics, Ulm University Hospital.

#### One Layer RDA

To assess the inhibitory effects of bacteriocin-producing bacteria, an one-layer RDA was performed, as previously described (Vogel et al., 2021). In short, the target bacteria were inoculated into trypticase soy agar (Oxoid) and poured into a plate. Subsequently, wells were placed in the agar and filled with the bacteriocin producing bacteria adjusted to an O.D. _600 nm_ of 0.5. After overnight incubation at 37 °C and 5% CO_2_, inhibition zones were measured in cm.

### Liposome leakage assay

The Folch method was used to extract lipids of live bacteria (*E. coli* and *P. aeruginosa*), as previously described (Folch et al., 1957; Vogel et al., 2022). Lipids were extracted from overnight cultures of bacteria. After harvesting, bacterial cells were resuspended in 1 ml of a 2:1(v/v) chloroform/methanol mixture and vortexed 5 x 1 min. To induce phase separation 200 µl of dH_2_O were added and centrifugation followed. The lipid phase was transferred into a glass vial. Liposome leakage assays were performed as previously described (Weil et al., 2020). Liposomes were prepared by thin-film hydration and extrusion. Lipids previously extracted or commercially purchased (*E. coli* polar,Avanti Polar Lipids) were hydrated with 50 mM 5(6)-carboxyfluorescein in 50% PBS (isoosmolar, pH 7.4), yielding a final lipid concentration of 5 mM. Samples were incubated at 70°C with shaking for 3 h, followed by 25 passes through 0.2 μm polycarbonate membranes using a Mini Extruder at 70°C. Free dye was removed by two rounds of size-exclusion chromatography (PD MidiTrap G-25), and liposome concentration was determined by nanoparticle tracking analysis (ZetaView).

For leakage assays, liposomes were diluted in PBS and 1–2 × 10^9^ particles were added per well in 96-well plates (80 μl). Fluorescence was monitored using a Synergy plate reader. After a 5 min baseline measurement, compounds (20 μl) were added and fluorescence recorded every minute for 1 h at 37°C. Complete dye release was induced with 2% Triton X-100. Data were background-subtracted and normalized to maximum fluorescence following detergent-induced lysis.

### AlphaFold structure prediction

ColabFold was used to predict the structure of Angicin and all its derivatives (Mirdita et al., 2022). In a second step the iCn3D viewer from NCBI, available at (https://www.ncbi.nlm.nih.gov/Structure/icn3d/), was used to view the structures.

### Prediction of good variants/ modification sites

Angicin variants were generated by replacing M9, E46, and M9+E46 for Lysine using the rational design tool from with CAMP_R4_ (Gawde et al., 2023) and then evaluated with CAMP_R4_-AMP prediction tool, Antimicrobial Peptide Scanner vr.2 (Veltri et al., 2018), and ABP-Finder (Ruiz-Blanco et al., 2022). Additionally, shorter AMP variants of 16-25 residues were generated with CAMP_R4_-Region tool and then evaluated with Antimicrobial Peptide Scanner vr.2 and ClassAMP (Joseph et al., 2012). A second group of shorter variants with molecular masses larger than 0.5 kDa were generated by *in silico* digestion of NGRWDYKVTKGIFQATTDVIVKGWA using Protein Digestion Simulator, https://pnnl-comp-mass-spec.github.io/Protein-Digestion-Simulator/ from PNNL and the Protein-Digestion-Simulator GitHub repository (accessed at 08.08.2018). The AMP prediction was done using CAMP_R4_ -AMP prediction tool, Antimicrobial Peptide Scanner vr.2, iAMPpred(Meher et al., 2017) and AI4AMP (Lin et al., 2021).

Last, from the active fragment KVTKGIFQATTDVIVKGWA two Cys-containing variants were proposed to form a cyclic peptide (A3.2.CC, N- and C-terminal Cys) and a homodimer (A3.2.C, N-terminal Cys) by disulfide bridges.

### *In vivo* toxicity assays in zebrafish embryos

*In vivo* toxicity was investigated using zebrafish embryos (*Danio rerio*). At 24 h post fertilization (hpf), wild-type embryos were dechorionated using digestion with 1 mg/mL pronase (Sigma) in E3 medium (83 μM NaCl, 2.8 μM KCl, 5.5 μM 202 CaCl2, 5.5 μM MgSO4). A total of 60 embryos were evaluated at each tested concentration, using two independent experiments of 10 groups of 3 embryos each. 3 embryos per well were exposed for 24 h to 100 µL of E3 containing 0.3, 3 and 30 µM Angicin. E3 medium was also used as a negative control. As a positive control for cytotoxicity 3 µM of the Pleurocidin antimicrobial peptide NRC-03 was used, cardiotoxicity was induced with 4% EtOH, and the neurotoxin Abamectin (3.125 µM) was administered as a positive control for neurotoxicity (Morash et al., 2011; Raftery et al., 2014). At 48 hpf (after 24 h of incubation), embryos were scored in a stereomicroscope for signs of cytotoxicity (lysis and/or necrosis, which is visible as loss of transparency), developmental toxicity (delay and/or malformations), or cardiotoxicity (heart edema and/or reduced or absent circulation). Each embryo was also touched with a needle, and a reduced, or absent touch response (escape movements) was evaluated as signs of neurotoxicity if and only if no signs of cytotoxicity were present in the same embryo. Embryos were categorized within each of these toxicity categories into several classes of severity as detailed in (Supplementary Table 6). Severe lysis (classes L4 and L3) and massive necrosis (Nec2) result in embryo death by the time of analyses, while embryos displaying the other phenotypic classes were still alive. Severely affected embryos (L4, L3, Nec2) were not analyzed for potential other phenotypes and were excluded from the graphs presenting data on developmental toxicity, cardiotoxicity and neurotoxicity. A Chi-Square test was used to calculate whether the distribution of embryos into toxicity classes differed significantly between treatment groups.

### Bioinformatic and Statistical analysis

Nucleotide sequences were retrieved from the GenBank database (http://www.ncbi.nlm.nih.gov/). The Basic Local Alignment Search Tool (http://www.ncbi.nlm.nih.gov/Blast/) was used to perform homology searches and to align protein sequences (Altschul et al., 1990). All other genetic analyses were performed with SnapGene 5.0 (https://www.snapgene.com/) and CLC Main Workbench v7.7.3 (http://www.clcbio.com). Statistical analyses was conducted with GraphPad Prism Version 10.5.0, GraphPad Software (http://www.graphpad.com).

## Author Contribution

VV and BS designed this study. AAS and LS, did bioinformatic analysis, predicted possible Angicin modifications and variants and synthesized them. BKN generated the *S. constellatus* mutant library and screened for Angicin-resistant mutants. LRO performed liposome leakage assays under the supervision of JM. JK conducted zebrafish toxicity experiments under the supervision of GW. PY developed a chemically defined medium for streptococci. JA prepared SEM samples with the help of CR and PW. All other experiments were performed by SM and VV. This manuscript was conceptualized and written by VV, JM and reviewed by BS.

## Aknowledgment

We thank Merve Balig and Nico Preising (Core Facility Functional Peptidomics) for their assistance in peptide synthesis.

## Data availability statement

The raw data supporting the conclusions of this article will be made available by the authors, without undue reservation.

## Ethics

Experiments were performed in accordance with German laws.

