## Supplementary Information for "Molecular Basis of Angicin Activity"

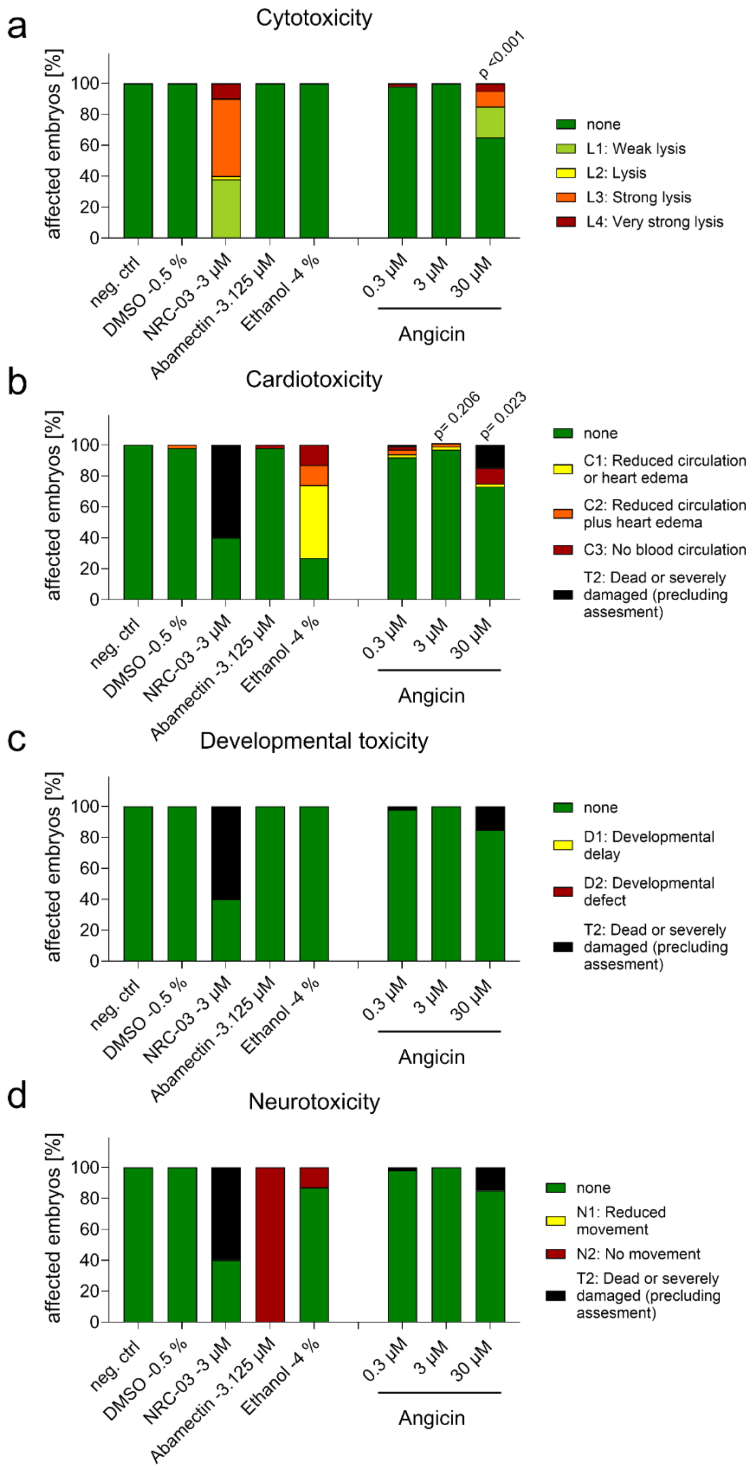

Supplementary Figure 1: Toxicity of Angicin in zebrafish embryos.

Zebrafish embryos (*Danio rerio*) were exposed to Angicin (0.3-30 µM) for 24h. Subsequently, cytotoxicity (a), cardiotoxicity (b), developmental (c) and neurotoxicity (d) were analysed with a stereomicroscope and touch response. In total, 60 embryos were exposed to each Angicin concentration in two independent experiments. As neg. ctrl, embryo medium was used. NRC-03, Abamectin and Ethanol serve as positive controls for cytotoxicity, neurotoxicity and cardiotoxicity, respectively. A Chi-square test was used to calculate significant differences.

Supplementary Table 1: Inactive peptides.

| Method | Two-layer RDA | | One-layer RDA |
|---|---|---|---|
| Antimicrobial active substance | AF3.2 | D-AF3.2 | BSU 1211 |
| Inactive against | *S. aureus, K. pneumoniae, S. anginosus* SK52*, S. constellatus* SK53, *E. coli, A. baumannii* | *S. aureus, K. pneumoniae, S. anginosus* SK52*, S. constellatus* SK53, *E. coli, P. aeruginosa, A. baumannii, E. faecium* (VRE), *L. monocytogenes* | *S. constellatus* SK53*manM*::ISS1 |
| Minimal amount of repetition | 2 | 2 | 2 |

Supplementary Table 2: Comparison of the protein sequences of Man-PTS transporter subunits IIC and IID.

Protein sequences were derived from publicly available nucleotide sequences (Accession numbers: *Streptococcus anginosus* SK52: LR134283; *Listeria monocytogenes* EGDe: AF397145; *Enterococcus faecium*: LR134337; *Escherichia coli*: CP154401; *Klebsiella pneumoniae*: CP167540) and compared to the protein sequence of *Streptococcus constellatus* SK53 (Accession number: *S. constellatus* SK53: AP014647). Depicted is percentage identity and query coverage.

| | **IIC**<br>**Identity/Query** | **IID**<br>**Identity/Query** |
|---|---|---|
| *S. anginosus* | 87.31 %/ 100 % | 86.47 %/ 100% |
| *L. monocytogenes* | 75.37 %/100 % | 67.77 %/ 100 % |
| *E. faecium* | 73.61 %/100 % | 68.77 %/ 99 % |
| *E. coli* | 46.64 %/99 % | 48.84 %/ 100 % |
| *K. pneumoniae* | 46.64 %/99 % | 47.52 %/ 100 % |

Supplementary Table 3: Bacteria used in this study.

| **Bacterial strain** | **Description** | **Source** |
|---|---|---|
| *Escherichia coli* BSU 1286 | ESBL, clinical isolate | University hospital Ulm |
| *Pseudomonas aeruginosa* BSU 856 | ATCC 27853 | ATCC |
| *Klebsiella quasipneumoniae* BSU 1353 | ESBL, ATCC 7000603 | ATCC |
| *Acinetobacter baumannii* BSU 1514 | *A. baumannii* type strain, ATCC 19606 | ATCC |
| *Staphylococcus aureus* BSU 1348 | MRSA, ATCC 43300 | ATCC |
| *Enterococcus faecium* BSU 1516 | VRE, DSM 17050 | DSMZ |
| *Streptococcus agalactiae* BSU 308 | ATCC 12403= NEM 316 | ATCC |
| *Streptococcus anginosus* SK 52 | *S. anginosus* type strain, ATCC 33397, Hly+ | ATCC |
| *Streptococcus anginosus* BSU 1211 | Clinical isolate | (Vogel et al., 2021) |
| *Listeria monocytogenes* EGDe BSU 1423 | Ln II Serotype I/2a | (Bécavin et al., 2014) |
| *Klebsiella pneumoniae* BSU 2231 | DSM 30104 | DSMZ |

Supplementary Table 4: Primers used in this study

| Primer Number | Name | Sequence 3'→ 5' |
|---|---|---|
| 1170 | manO_rev | gtatcacggtagaatttcaag |
| 1171 | manM_rev | catccgttcatagttccaag |
| 1172 | ManX_fwd | gtttacaatctgtcaaaaatatg |
| 1173 | ManN_fwd | ggagctactaatatcatgacag |
| 1205 | ManXM_fwd | agcatcaggcgcaacagcag |
| 1249 | ManX_end_fwd | gccaacgttcaaataatagaaagg |

Supplementary Table 5: Cytotoxicity classification for zebrafish embryos.

| **Cytotoxicity** | |
|---|---|
| L1: | few lysed cells floating in medium; embryos look like wt |
| L2: | lysed cells in medium; embryos show some visible tissue damage |
| L3: | embryos show strong tissue damage, they are so severely damaged that assessment of all other types of toxicity does not make sense |
| L4: | embryos are completely disintegrated |
| Nec1: | individual necrotic cells (darkened areas in brightfield) |
| Nec2: | many necrotic cells (darkened areas in brightfield) |
| **Developmental toxicity** | |
| D1: | developmental delay (slow development) |
| D2: | developmental defect (malformations) |
| **Cardiotoxicity** | |
| C1: | reduced circulation or heart edema |
| C2: | reduced circulation plus heart edema |
| C3: | no circulation with or without heart edema |
| **Neurotoxicity** | |
| N0: | normal movement in response to touch |
| N1: | reduced movement in response to touch |
| N2: | no movement in response to touch |
| **Overall toxicity (combination of above phenotypes)** | |
| wt: | wild type (no visible phenotype AND normal movement) |
| T1: | embryos that show a phenotype |
| T2: | severe damage so that other phenotypes cannot be assessed (L3, L4, Nec2) |

### Supplementary Methods

#### Preparation of chemically defined medium for streptococci

Firstly, the following Stock solutions were prepared.

25 x Phosphate Stock solved in $dH_2O$

| Amount in g/l | Component |
|---|---|
| 5 | $K_2HPO_4$ |
| 25 | $KH_2PO_4$ |
| 79.9 | $NaH_2PO_4$ -$H_2O$ |
| 346.8 | $Na_2HPO_4$ -7 $H_2O$ |

500 x Magnesium Sulfate Stock in $dH_2O$

350 g/l of $MgSO_4$ -7 $H_2O$

1000 x Manganese (II) sulfate Stock in $dH_2O$

2.5 g/l $MnSO_4$

50 x Sodium Acetate Stock in $dH_2O$

225 g/l $NaC_2H_3O_2$-3 $H_2O$

20 x Sodium Bicarbonate Stock in $dH_2O$

65 g/l NaHCO3

1000 x Calcium Chloride Stock in $dH_2O$

10 g/l $CaCl_2$-2 $H_2O$

1000 x Iron Stock in $dH_2O$

1 g/l $Fe(NO_3)_3$-9$H_2O$

5 g/l $FeSO_4$-7 $H_2O$

500 x L-Cysteine Hydrochloride Stock in $dH_2O$

250 mg/l $C_3H_7NO_2S$-HCl

100 x Base Stock in 2.4 M HCl

2 g/l adenine

2 g/l guanine hydrochloride

2 g/l uracil

1000 x Vitamins (Vitamins Kit, Sigma) Stock in $dH_2O$

| Amount in g per l | Component |
|---|---|
| 0.2 | p-aminobenzoic acid |

| | |
|---|---|
| 0.2 | biotin |
| 0.8 | folic acid |
| 1 | nicotinamide |
| 2 | pantothenate calcium salt |
| 1 | pyridoxal |
| 1 | pyridoxamine dihydrochloride |
| 2 | riboflavin |
| 1 | thiamine hydrochloride |
| 0.1 | vitamin B12 |

50 x Amino Acid Stock in $dH_2O$

| Amount in g per l | Amino acid |
|---|---|
| 5 | L-alanine |
| 5 | L-arginine |
| 5 | L-aspartic acid |
| 2.5 | Thymine |
| 5 | L-cystine |
| 10 | L-glutamic acid |
| 5 | Glycine |
| 5 | L-histidine |
| 5 | L-isoleucine |
| 5 | L-leucin |
| 5 | L-lysine |
| 5 | L-methionine |
| 5 | L-phenylalanine |
| 5 | L-proline |
| 5 | Hydroxy-L-proline |
| 5 | L-serine |
| 10 | L-threonine |
| 5 | L-tryptophan |
| 5 | L-tyrosin |
| 5 | L-valin |

For CDM the Stock solutions were mixed according to Table 1.

Table 1: Recipe for CDM

| Amount in ml | Component |
|---|---|
| 40 | 25 x Phosphate Stock |
| 2 | 500 x Magnesium Sulfate Stock |
| 1 | 1000 x Manganese (II) sulfate Stock |
| 20 | 50 x Sodium Acetate Stock |
| 50 | 20 x Sodium Bicarbonate Stock |
| 1 | 1000 x Calcium Chloride Stock |

| | |
|---|---|
| 1 | 1000 x Iron Stock |
| 2 | 500 x L-Cysteine Hydrochloride Stock |
| 1 | 1000 x Vitamins Stock |
| 10 | 100 x Base Stock |
| 20 | 50 x Amino Acid Stock |
| Variable | Carbon source (e.g. D-Glucose, D-Mannose) |
| Ad 1 l d$H_2O$ | |

All Components were mixed and sterile filtered. CDM was prepared freshly every two weeks.
